# Bacterial diadenylate cyclase domains synthesize diverse nucleotide signals in anti-phage defense

**DOI:** 10.64898/2026.08.10.743927

**Authors:** Adelyn E. Ragucci, Danielle J. Haley, Philip J. Kranzusch

## Abstract

Bacterial diadenylate cyclase (DAC) enzymes synthesize the nucleotide signal 3′3′ cyclic di-AMP (3′3′-c-di-AMP) to control osmoregulation, cell-wall homeostasis, and DNA-damage responses. Here we discover specialized roles for DAC enzymes in bacterial immunity and define a Panoptes-like system we name Panoptoo as a DAC-containing anti-phage defense that guards against viral immune evasion. The Panoptoo protein PanS is a minimal DAC that constitutively synthesizes 3′3′ cyclic UMP-AMP (3′3′-cUA) or 3′3′-c-di-AMP to negatively regulate a partnering PanE S2TMβ membrane-targeting effector. We show that Panoptoo decoy signaling acts as a counter-defense to detect phage immune evasion proteins that inhibit nucleotide immune signals. A 1.5 Å crystal structure of PanS in complex with 3′3′-cUA explains how a symmetry break in the canonical DAC active site enables synthesis of asymmetric signaling molecules. Together, our results uncover a role for DAC domains in bacterial anti-phage defense and expand our understanding of nucleotide signaling in antiviral immunity.

## Introduction

Nucleotide second messenger signaling pathways are a major form of antiviral immunity conserved across all kingdoms of life^1–3^. Especially prevalent forms of nucleotide immune signaling include CBASS (cyclic oligonucleotide based anti-phage signaling system), Pycsar (pyrimidine cyclase system for anti-phage resistance), Thoeris, Kongming, and type III CRISPR anti-phage defense systems in bacteria^4–11^, TIR domain immune effector proteins in plants^7,8,12,13^, and cGAS (cyclic GMP-AMP synthase) and other cGLR (cGAS-like receptor) signaling pathways in human and animal antiviral immunity^14–17^. In each case, recognition of viral infection induces enzymatic synthesis of specialized nucleotide immune signals that then bind to downstream receptors to control immune effector responses^2,3^. In addition to pathways that synthesize nucleotide signals as positive regulators of immune activation, recent discoveries of Hailong, Panoptes, and Clover anti-phage defense systems in bacteria demonstrate that nucleotide immune signals can function as negative regulators that repress immune activation until recognition of viral infection^18–21^. Together, nucleotide immune signaling pathways enable coordinated regulation of immune responses and constitute a fundamental mechanism of how cells sense and defend against viral infection.

Competition between viruses and cells drives evolution and extensive diversification of nucleotide immune signaling pathways. Viruses encode immune evasion proteins that function as enzymes to degrade nucleotide immune signals or sponges that bind and tightly sequester signals to disrupt immune activation^2,3,6,22–29^. In response, host cells use alteration of nucleobase specificity, distinct phosphodiester linkage chemistries, and incorporation of additional metabolite classes as mechanisms to adapt nucleotide signals and restore antiviral immunity^4–8,13,15–17,30–33^. Nearly all major protein domains in biology demonstrated to synthesize nucleotide signals are known to have roles in antiviral defense including mcPol / GGDEF / adenylate cyclase ferredoxin-like fold enzymes in type III CRISPR, Panoptes, and Pycsar defense^6,10,11,19,20^; CD-NTase and cGLR enzymes in CBASS defense and animal innate immunity^4,5,14,17^; and TIR domains in bacterial, plant, and animal immunity^7,8,13,33,34^. A notable exception are diadenylate cyclase (DAC) domain proteins that are especially prevalent signaling enzymes in gram-positive bacteria that synthesize the nucleotide signal 3′3′ cyclic di-AMP (3′3′-c-di-AMP) to control osmoregulation and the response to DNA damage^35,36^. DAC domain proteins have been bioinformatically identified in bacterial anti-phage defenses^4,37–40^, suggesting that these enzymes may also function as nucleotide immune signal generating enzymes in antiviral immunity.

Here we biochemically and structurally characterize DAC domain proteins in bacterial immunity and demonstrate that these enzymes synthesize specialized nucleotide signals to control anti-phage defense. We discover Panoptoo as a two-gene operon (*panSE*) that guards against phage immune evasion proteins through negative regulation of the PanE S2TMβ effector. Surprisingly, we find that most PanS diadenylate cyclase enzymes constitutively synthesize 3′3′ cyclic UMP-AMP (3′3′-cUA), a nucleotide product distinct from the canonical DAC domain product 3′3′-c-di-AMP. Structural analysis of the PanS post-reactive state reveals two distinct protomer conformations that explain how the dimeric PanS DAC domain has been adapted to synthesize an asymmetric signaling molecule. Our results explain how bacteria use DAC domains to expand antiviral immunity and add to the growing repertoire of nucleotide signals that negatively-regulate immune activation.

## Results

### DAC domain proteins are signaling enzymes in bacterial defense islands

To identify signaling enzymes in bacterial immunity, we analyzed anti-phage defense islands for proteins with PFAM signatures associated with nucleotide second messenger synthesis and identified two distinct classes of proteins containing the DAC domain known to synthesize the cyclic dinucleotide signal 3′3′c-di-AMP^35,41^ (Figure 1a,b). The first class of defense-associated DAC proteins are large ~550 amino acid proteins previously found to occur within a variant of CBASS operons designated DS-2^4,37–40^ (Extended Data Figure 1a). These proteins, referred to as DS-2C, are predicted to contain three domains: DACNG (InterPro IPR048554) and DACNH (InterPro IPR048555), two uncharacterized domains located at the N-terminus of bacterial DACs and hypothesized to function as sensory modules^37^, and a canonical DAC DisA_N domain with predicted similarity to the core enzymatic domain of the bacterial DNA integrity scanning protein that synthesizes 3′3′-c-di-AMP^35,41^ (Extended Data Figure 1b,c). The second class of defense-associated DAC proteins are minimal ~150 amino acid proteins also recently identified in large-language model prediction methods for unbiased annotation of bacterial immunity proteins^38,39^. These smaller proteins contain only the canonical DAC DisA_N domain and occur as part of two-gene operons with a partnering two transmembrane β-strand rich (S2TMβ) protein that has homology to the nucleotide signaling receptor Cap15 in CBASS immunity^42^ (Extended Data Figure 1a). The individual DAC domains from each defense-associated protein are divergent and exhibit only ~27% and ~25% amino-acid identity with *B. subtilis* DisA. To enable comparison, we therefore modeled structures of representatives from each class using AlphaFold3^43^ and observed that the structural models for the predicted defense-associated DAC proteins retain each of the architectural elements known to be required for DisA enzymatic activity including a central four-stranded β-sheet β1–β4 braced by conserved helices α1–α5 (Figure 1a,b and Extended Data Figure 1c). DALI^44^ analysis of the DS-2C DAC domain and DisA_N protein AlphaFold3 models compared to structures in the Protein Data Bank confirmed that these defense-associated proteins share strong homology to DisA and CdaA diadenylate cyclases^35,45,46^ (Figure 1c, Extended Data Figure 1d). Furthermore, we used the AlphaFold3 models to guide sequence-alignment of the divergent DAC domains and observed conservation of select residues within the core active site motifs DGA and RHR required for substrate recognition and nucleotide second messenger formation, suggesting that each defense-associated protein is an active signaling enzyme^45,47^ (Extended Data Figure 1e,f).

**Figure 1.**
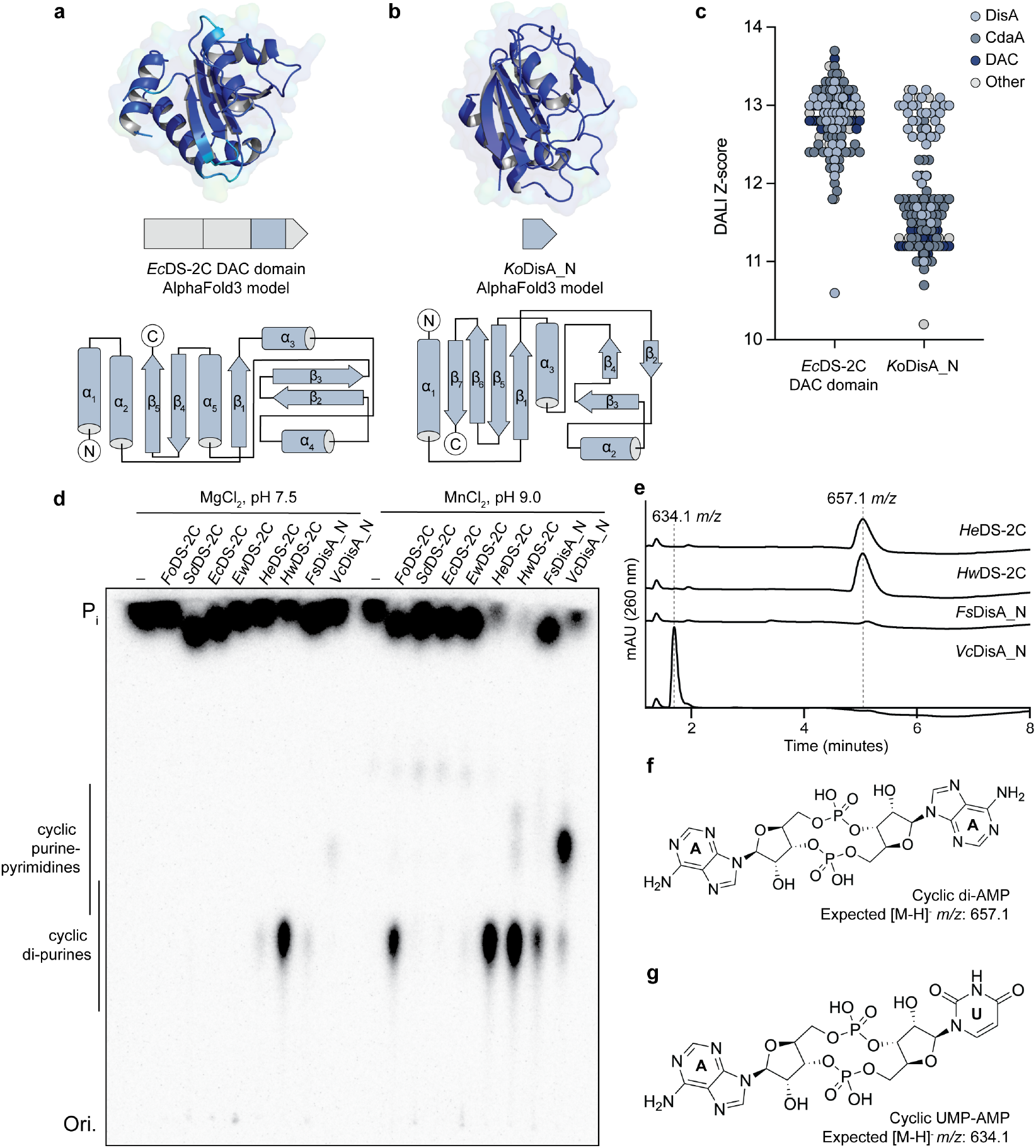
Defense-associated DAC domains synthesize diverse cyclic dinucleotides. **a,b**, AlphaFold3 predicted structures of the *Ec*DS-2C (WP_001593457.1) DAC and *Ko*DisA_N (WP_071889451.1) DAC domain structures colored by pLDDT. Topology diagrams are depicted below. **c**, Z-score structural similarity plot demonstrating homology between *Ec*DS-2C DAC domain and *Ko*DisA_N model predictions with representative structures in the Protein Data Bank. Z-score cut-offs for both models are 10. **d**, Thin-layer chromatography (TLC) analysis of radiolabeled biochemistry reactions for purified DS-2C and DisA_N incubated with all four NTPs at pH 7.5 with MgCl2 or pH 9.0 with MnCl_2_ demonstrates that defense-associated DAC domains synthesize cyclic di-purine and cyclic purine-pyrimidine containing nucleotide products. **e**, LC-MS analysis of reactions performed at pH 9.0 with MgCl_2_ and MnCl_2_ verifies that DS-2C and *Fs*DisA_N synthesize c-di-AMP and *Vc*DisA_N synthesizes cUA. **f,g**, Example chemical structures of c-di-AMP and cUA with canonical 3′–5′ phosphodiester bonds. Data shown in **d–e** are representative of three independent experiments.

We selected defense-associated DAC domain-containing proteins representing each class and purified eight recombinant proteins to test for signaling activity *in vitro* (Figure 1d, Extended data Figure 1g). Using a range of reaction conditions testing substrate (adenosine triphosphate (ATP), guanosine triphosphate (GTP), cytidine triphosphate (CTP), uridine triphosphate (UTP)), metal co-factor (Mg^2+^, Mn^2+^), and pH (7.5, 9.0) preferences, we observed robust nucleotide second messenger synthesis activity for five proteins: *Flagellimonas okinawensis* DS-2C, *Hymenobacter elongatus* DS-2C, *Hymenobacter wooponensis* DS-2C, *Flavihumibacter sp. UBA7668* DisA_N, and *Vibrio coralliirubri* DisA_N (Figure 1d). Notably, DAC domain-containing proteins synthesized at least two different products with *H. wooponensis* DS-2C and *Flavihumibacter sp*. DisA_N synthesizing a product that migrated on polyethyleneimine (PEI) cellulose thin-layer chromatography (TLC) consistent with a cyclic dipurine molecule and *V. coralliirubri* DisA_N synthesizing a product with a distinct TLC migration indicative of a cyclic purine–pyrimidine molecule^4,17^ (Figure 1d). To further interrogate the identity of the nucleotide products, we analyzed reactions with liquid chromatography mass spectrometry (LC-MS) and observed that *H. elongatus* DS-2C, *H. wooponensis* DS-2C, and *Flavihumibacter sp*. DisA_N reactions produced a peak species that corresponds to a c-di-AMP molecule in agreement with the canonical DAC product 3′3′-c-di-AMP (Figure 1e,f). In contrast, the *V. coralliirubri* DisA_N reaction product migrated as a unique peak with a retention time and mass corresponding to a c-UMP-AMP molecule (cUA) (Figure 1e,g). These results demonstrate that DAC domains associated with anti-phage defense islands are functional nucleotide second messenger synthesis enzymes that produce diverse signals.

### Panoptoo PanS synthesizes cUA and c-di-AMP as negative regulators of operon signaling

The ability of defense-associated DisA_N proteins to synthesize cUA is distinct from all previously characterized DAC domains that produce the signal c-di-AMP^35,41^, and we therefore decided to focus characterization on operons containing these minimal DAC proteins. We cloned full-length operons of the two-gene DisA_N and partnering S2TMβ protein systems from *Proteobacteria* species and expressed each construct in *E. coli* using an arabinose-inducible promoter^30,48^. Although expression of the full-length operons was tolerated, we observed that expression of the S2TMβ proteins alone induced potent cell toxicity, suggesting that the systems may require DisA_N nucleotide second messenger synthesis to negatively regulate effector function (Figure 2a). These results are reminiscent of a recently characterized form of anti-phage defense named Panoptes that uses a minimal CRISPR polymerase (mcPol)-domain containing protein to synthesize 2′3′-c-di-AMP or cyclic tri-AMP and negatively regulate activation of an S2TMβ or CARF-TM effector^19,20,49^ (Extended Data Figure 2a). We therefore named the DisA_N protein containing operons “Panoptoo” as a two-gene system *panSE* encoding the nucleotide synthase PanS and the partnering effector protein PanE. We compared Panoptes and Panoptoo genomic neighborhoods and observed key shared traits (Figure 2b). First, each system is composed of a two-gene unit where the nucleotide second messenger signaling protein (OptS or PanS) is near universally encoded next to an S2TMβ effector protein (OptE or PanE). Second, both systems are most frequently encoded adjacent to CBASS operons allowing negative regulation in Panoptes and Panoptoo signaling to guard CBASS systems against phage immune evasion proteins^19,20^. These results support Panoptes and Panoptoo as sister systems where alternative replacement of the core nucleotide second messenger synthesis domain with either mcPol- or DAC-containing proteins allows diversification of nucleotide immune signaling.

**Figure 2.**
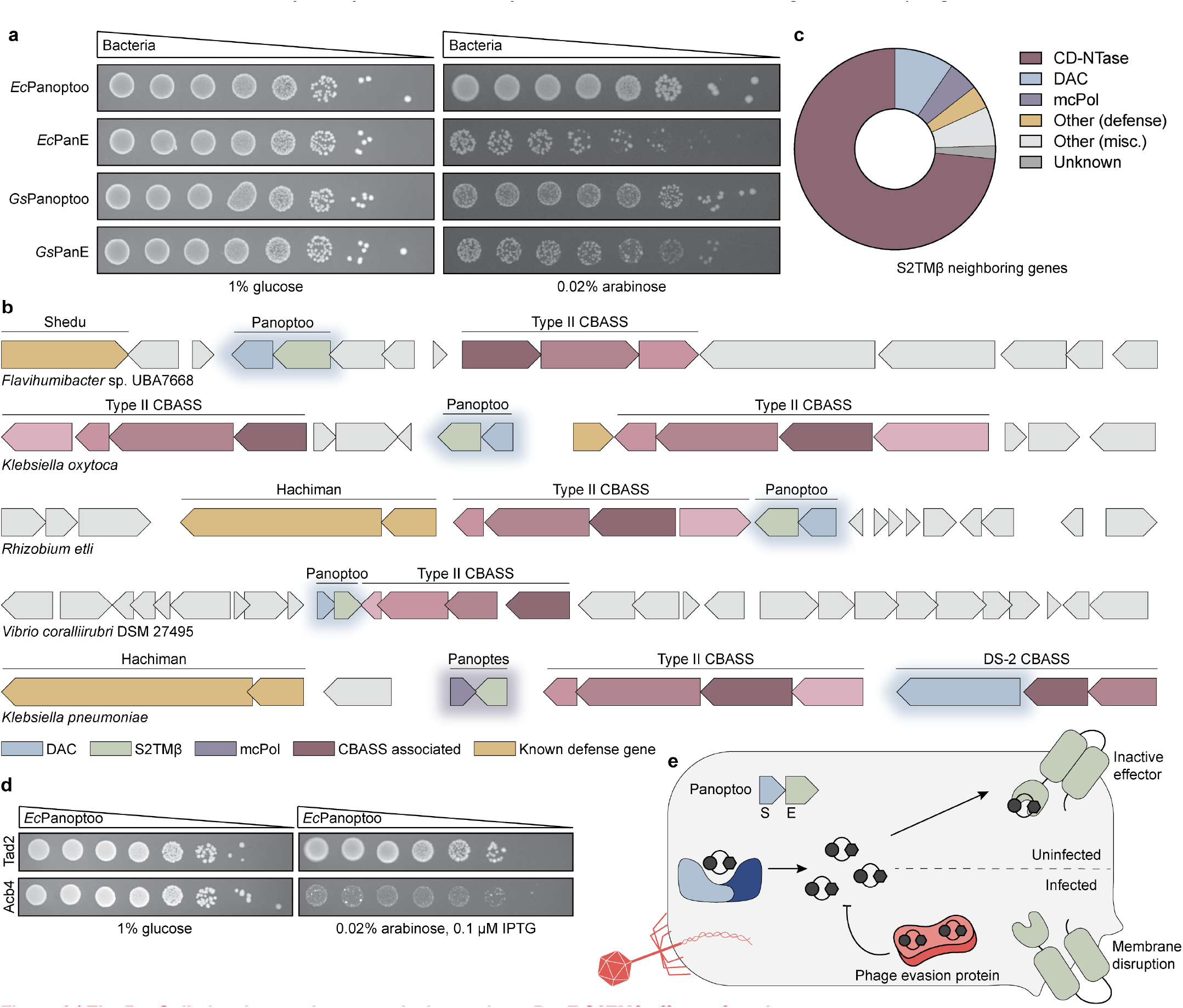
The PanS diadenylate cyclase negatively regulates PanE S2TMβ effector function. **a**, Bacterial growth spot assay of *E. coli* expressing a complete Panoptoo PanSE system or the PanE effector alone. Protein expression is under the control of an arabinose inducible promoter. Bacteria grown in repressive (1% glucose) or inducing (0.02% arabinose) conditions reveal that expression of PanE alone induces cellular toxicity. **b**, Gene neighborhood analysis for representative Panoptoo and Panoptes defense operons. **c**, Analysis of the distribution of neighboring genes for 617 S2TMβ effectors demonstrates that S2TMβ effectors most frequently occur as part of CBASS (73.4%), Panoptoo (9.6%), and Panoptes (4.9%) anti-phage defense systems. **d**, Bacterial growth assay analyzing co-expression of *Ec*Panoptoo with phage SPO1 Tad2 and Acb4 immune evasion proteins. Co-expression of *Ec*Panoptoo and SPO1 Acb4 specifically induces cellular toxicity in agreement with the known ability of SPO1 Acb4 to sequester 3′3′-cUA nucleotide immune signals^27^. **e**, Model depicting the mechanism of Panoptoo defense in uninfected and phage infected states. PanS constitutively synthesizes a nucleotide signal that negatively regulates PanE. Phage immune evasion proteins that disrupt nucleotide immune signaling induce activation of PanE and membrane disruption. Data shown in **a** and **d** are representative of three independent experiments.

To further understand diversity of this signaling architecture, we constructed a phylogenetic tree of S2TMβ effector sequences (InterPro IPR041208) and analyzed each protein for co-occurrence with potential nucleotide signal generating enzymes (Extended Data Figure 2c). In analysis of 617 S2TMβ effectors, 73.4% are Cap15 effectors associated with a CD-NTase enzyme as part of CBASS anti-phage defense, 9.6% are PanE effectors in Panoptoo systems, and 4.9% are OptE effectors in Panoptes systems (Figure 2c). The remaining S2TMβ effectors are often encoded in defense islands next to a variety of proteins including α/β fold hydrolases, helix-turn-helix proteins, and tyrosine recombinases that may represent additional functional operons (Extended Data Figure 2e). To investigate whether diadenylate cyclases are encoded as guards in bacterial immunity, we co-expressed the full-length Panoptoo operon with phage immune evasion proteins. The phage SPO1 proteins Tad2 and Acb4 are known to bind the signaling molecules produced from Thoeris (3′-cADPR and 2′-cADPR) and CBASS (3′3′-cGAMP, 2′3′-cGAMP, 3′2′-cGAMP, 3′3′-cUA and 2′3′-cUA) anti-phage defense, respectively^26,27,33^. Phage SPO1 Tad2 is unable to bind cUA-like signaling molecules^26,33^ and as expected, no toxicity was observed upon co-expression of the *E. coli* ECC-Z Panoptoo operon with SPO1 Tad2 (Figure 2d). However, phage SPO1 Acb4 binds and sequesters cUA with high-affinity^27^ and we observed robust cell toxicity upon co-expression of the *E. coli* ECCZ Panoptoo operon with SPO1 Acb4 demonstrating loss of negative regulation and activation of anti-phage defense (Figure 2d). Together, these results establish Panoptoo as a guard-based defense system that uses negative regulation of S2TMβ effectors to subvert phage anti-CBASS immunity (Figure 2e).

### The primary signal in Panoptoo defense is 3′3′-cUA

To begin to understand the molecular basis of Panoptoo nucleotide immune signaling, we generated a phylogenetic tree of 200 PanS homologs encoded next to S2TMβ effectors and selected 9 additional homologs to express, purify, and biochemically characterize (Figure 3a, Extended Data Figure 3a). Upon expression and purification of the 11 PanS proteins in our screen, we found that each protein exhibited a high A_260_ absorbance and co-purified with a bound nucleotide signaling molecule consistent with constitutive signaling in guard-like anti-phage defense^18–21^ (Extended Data Figure 3b,c). We denatured the PanS proteins to release the bound signaling molecules for LC-MS analysis and observed that 9 of the 11 tested PanS homologs co-purified with cUA and 2 of the PanS homologs co-purified with c-di-AMP (Figure 3b). We next reconstituted PanS signaling for each homolog *in vitro* with radiolabeled nucleotide substrates and confirmed cUA synthesis for PanS homologs from *Aeromonas salmonicida, Pseudomonas marginalis, Rhizobium etli, Shewanella algae, Variovorax paradoxus*, and *Wohlfahrtiimonas chitiniclastica* bacterial species and c-di-AMP synthesis for PanS homologs from *Flavihumibacter sp*. and *Giesbergeria sp*. (Figure 3c, Extended Data Figure 3d). For one PanS homolog from *V. coralliirubri* we observed cUA as a major product and c-di-AMP as a minor product, suggesting that individual PanS systems may adapt nucleotide immune signal identity similar to known variability in CBASS, Pycsar, and type III CRISPR nucleotide signaling systems^4,6,10,11^ (Figure 1d, Figure 3c). Finally, to determine the phosphodiester linkage specificity of the nucleotide immune signals in Panoptoo defense we combined nucleo-base-specific labeling of the PanS cUA and c-di-AMP products with nuclease P1 digestion that specifically degrades 3′–5′ phosphodiester bonds and is unable to degrade 2′–5′ bonds^4,15,50^. We observed complete digestion with nuclease P1 confirming the PanS products as 3′3′-cUA and 3′3′-c-di-AMP (Figure 3d,e,f). These results demonstrate that PanS proteins constitutively synthesize diverse cyclic dinucleotide signals with 3′3′-cUA as the predominant signaling molecule in Panoptoo defense.

**Figure 3.**
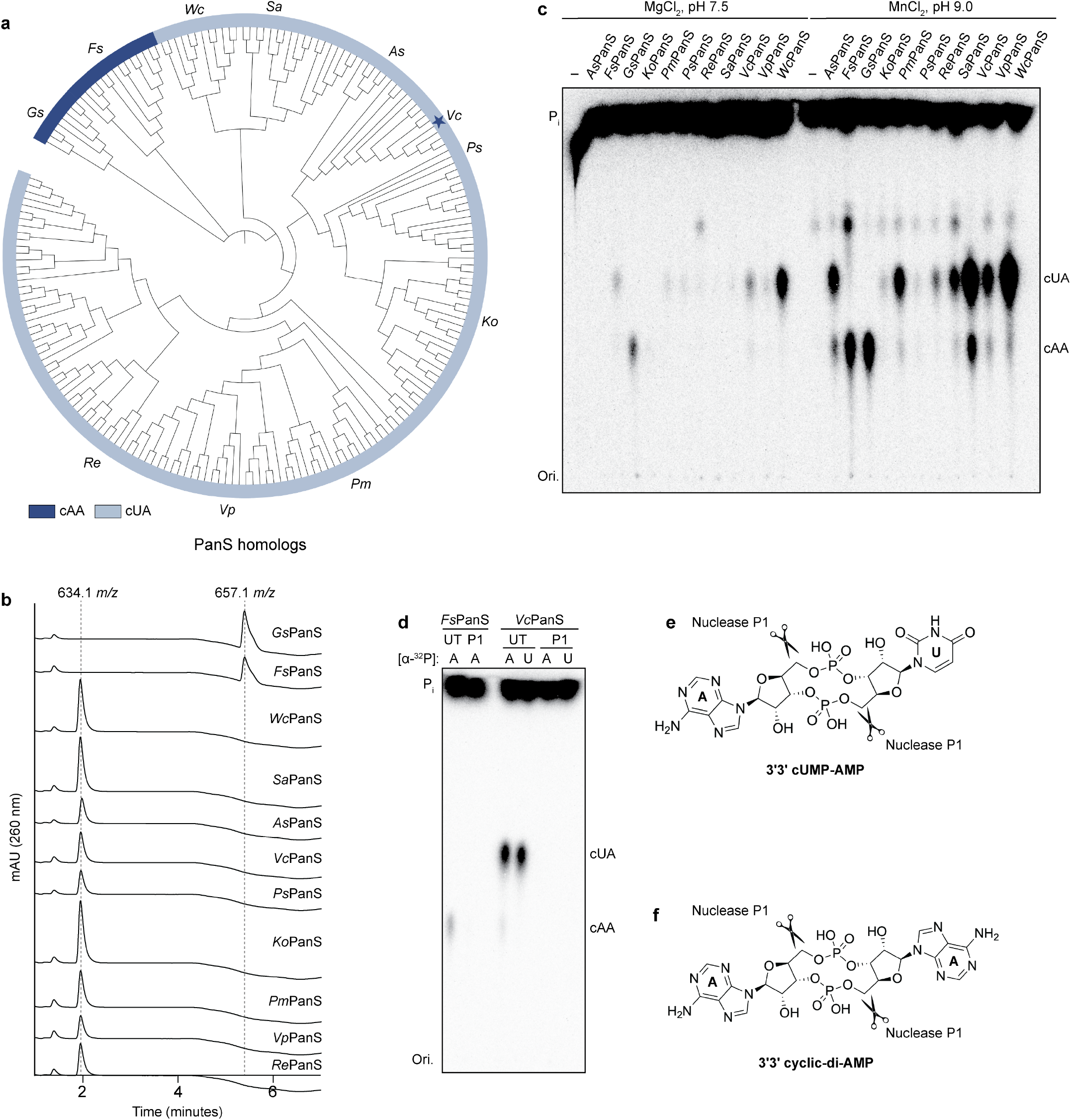
PanS enzymes synthesize 3′3′-c-di-AMP or 3′3′-cUA. **a**, Phylogenic analysis of 200 PanS homologs encoded next to PanE effectors. Enzymes experimentally tested are annotated on the outside of the tree enabling definition of PanS homologs that synthesize cyclic di-AMP (cAA, dark blue) and cyclic UMP-AMP (cUA, light blue). *Vc*PanS synthesizes cUA as a major product and cAA as a minor product and is denoted with a blue star. **b**, LC-MS analysis of 11 PanS enzymes demonstrates copurification of cUA and cAA nucleotide signals. **c**, TLC analysis of radiolabeled biochemistry reactions for 11 purified PanS enzymes confirms that most PanS enzymes synthesize cUA but select species including *Fs*PanS and *Gs*PanS synthesize cAA. **d**, TLC analysis of *Fs*PanS and *Vc*PanS reaction products treated with nuclease P1 confirms that PanS homologs synthesize nucleotide signals with canonical 3′–5′. UT represents untreated samples while P1 indicates nuclease P1 treated samples. **e–f**, Chemical structures of the PanS nucleotide signals 3′3′-cUA (**e**) and 3′3′-c-di-AMP (**f**) annotated with cleavage sites for nuclease P1. Data shown in **c–d** are representative of three independent experiments.

### PanS active site remodeling enables synthesis of asymmetric ligands

To define the mechanism of 3′3′-cUA synthesis in Panoptoo defense, we next determined a 1.5 Å crystal structure of PanS from the bacterium *R. etli* (Figure 4a, Extended Data Figure 4a; Extended Data Table 1). The PanS structure reveals a post-reaction state with three sets of PanS dimers each embraced around a molecule of 3′3′-cUA in the enzyme active site (Figure 4a,b, Extended Data Figure 4b). The active PanS dimer adopts an overall architectural conformation conserved with canonical DAC enzymes with PanS and *Thermotoga maritima* DisA exhibiting an RMSD of 3.1 Å^35^. In previous structures of DisA, the core enzyme dimer forms a perfectly two-fold symmetric active site in-line with synthesis of the symmetric product 3′3′-c-diAMP^35,45^. In stark contrast, the PanS structure reveals two distinct protomer conformations that remodel the enzyme and create an asymmetric active site (Figure 4c–f; Extended Data Figure 4c). Analysis of PanS and DisA nucleotide contact maps explain how rearrangement of residues forming the “lid” of PanS breaks the canonical two-fold symmetric axis and enables PanS to specifically direct 3′3′-cUA synthesis (Figure 4g,h). For both DisA and PanS, residues of the DGA and RHR active site motifs make conserved contacts to the phosphate and ribose backbone of cyclic di-AMP and 3′3′-cUA, respectively^35^ (Figure 4g,h, Extended Data Figure 4d–g). In both protomer conformations of PanS, the position of residues D71, G72, G107, and R108 at the bottom of the active site remains the same, enabling the enzymatic reaction necessary for concerted phosphodiester bond formation (Figure 4e–g,j). However, large conformational changes in a loop region between A86 and I93 in the enzyme lid remodels the top of the PanS active site into distinct conformations that allows the enzyme to accommodate adenine or uracil bases (Figure 4c–g). In particular, S90 is pointed away from the adenine pocket in the first PanS protomer conformation but undergoes a 4 Å shift such that the side chain points down toward the uracil pocket in the second protomer conformation (Figure 4c–g). This conformational change effectively closes the PanS lid and remodels the enzyme for synthesis of an asymmetric signaling molecule. Compared to PanS, this loop region is shorter and more structured in DisA, CdaA, and CdaS^35,45,46,51^, explaining why canonical DAC proteins don’t undergo similar structural rearrangement (Figure 4i). Together, these data demonstrate how PanS proteins remodel a flexible lid region in DAC domains to enable asymmetric nucleotide product synthesis and function in a specialized role in anti-phage defense.

**Figure 4.**
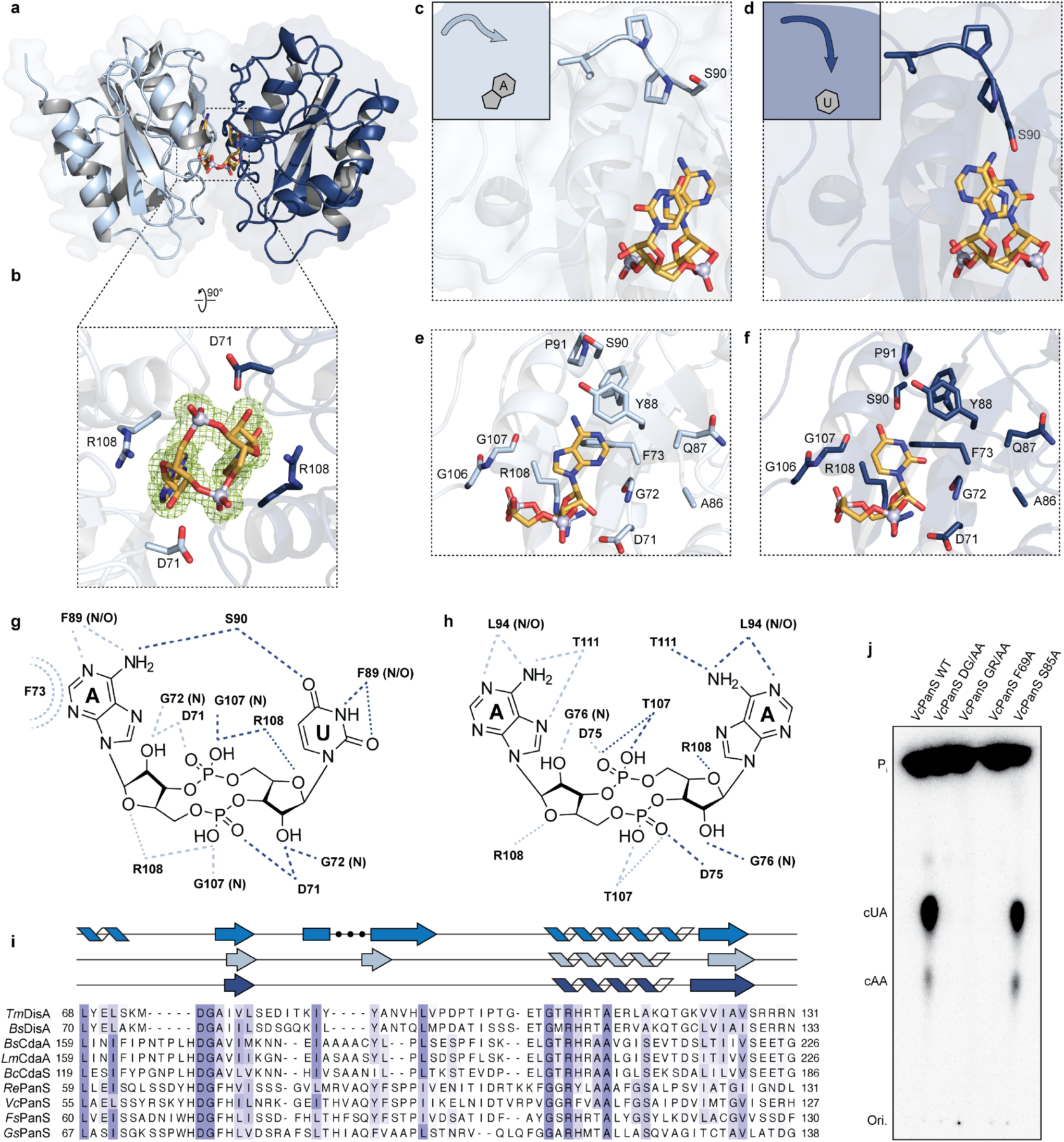
Mechanism of PanS 3′3′-cUA product formation. **a**, Overview of the crystal structure of *Re*PanS in complex with the product nucleotide signal 3′3′-cUA. **b**, Zoomed view of the *Re*PanS active site with 3′3′-cUA ligand and conserved catalytic contacts shown as sticks. The bottom half of the *Re*PanS active site is symmetric, where D71 and R108 read out the phosphodiester linkage of 3′3′-cUA. Green mesh depicts F_O_–F_C_ polder omit map (σ = 3). **c–d**, Cartoon and zoomed-in schematic of the two conformations of the *Re*PanS loop lid region that controls nucleotide specificity. In the protomer which contacts adenosine (light blue), S90 is turned away from the binding pocket (**c**). In the protomer which contacts uracil (dark blue), S90 is turned toward the binding pocket and makes contacts with both nucleobases (**d**). **e–f**, Zoomed in cutaway of the 3′3′-cUA binding pocket with ligand-contacting and loop residues shown as sticks. **g–h**, Contact maps highlighting ligand recognition in *Re*PanS (**g**) compared to the canonical DAC protein *Tm*DisA (**h**). The active site of *Tm*DisA is completely symmetric. In contrast, the *Re*PanS active site is asymmetric with only the contacts made at the bottom of the *Re*PanS active site are shared between protomers. Hydrogen bonding is depicted as dashed lines while water bridges are shown as dotted lines. **i**, Sequence alignment of DAC proteins demonstrates that the flexible extension in the PanS lid region is absent in canonical DisA, CdaA, and CdaS enzymes. Structure guided sequence alignment (top) demonstrates that this region is structured in *Tm*DisA (bright blue) while the loop has two different conformations in the *Re*PanS protomers (light and dark blue). **j**, Mutants in the conserved catalytic sites (DGF and GGR) of *Vc*PanS result in loss of 3′3′-cUA production while S85 is dispensable for enzymatic activity. Data are representative of three independent experiments.

## Discussion

Here we define a role for diadenylate cyclases in bacterial immunity and discover that DAC domain-containing proteins synthesize diverse nucleotide signals to control anti-phage defense. Building on the recent discovery of Panoptes defense^19,20^, we define Panoptoo as a related system where PanS synthases are DAC enzymes that create decoy signals including 3′3′-cUA and 3′3′-c-di-AMP to indirectly sense phage immune evasion proteins. We show that in Panoptoo defense, disruption of PanS signaling releases activity of the membrane-targeting antiviral effector PanE (Figures 1 and 2). We additionally analyze DS-2C DAC proteins associated with CBASS operons^37^ and demonstrate that these larger DAC protein variants are active *in vitro* and synthesize the signal 3′3′-c-di-AMP (Figure 1). Together our results reveal that bacterial immune systems expand both the roles and nucleotide specificity of DAC domain protein function.

PanS synthesis of 3′3′-cUA reveals a unique ability for a DAC protein to create an asymmetric nucleotide signal (Figures 1 and 3). All previously characterized DAC proteins form symmetric, multimeric assemblies that catalyze synthesis of the 2-fold symmetric signal 3′3′-c-di-AMP^35,41^. 3′3′-c-di-AMP is a cellular homeostasis signal typically used for monitoring DNA damage, cell wall osmoregulation, and other stress responses in bacteria of the *Firmicutes, Actinobacteria*, and *Cyanobacteria* phyla^36^. In contrast, we show that diverse *Proteobacteria* species encode PanS homologs as DAC proteins that synthesize the noncanonical product 3′3′-cUA dedicated to anti-phage defense (Figures 3 and 4). Structural analysis of the PanS postreaction state explains how distinct conformations in the enzyme flexible lid region create a symmetry break in the active-site that alternatively accommodates either adenine and uracil bases to complete 3′3′-cUA product formation (Figures 3 and 4). We show that in some Panoptoo systems the PanS proteins synthesize the canonical DAC domain product 3′3′-c-di-AMP, suggesting that alteration in nucleotide base specificity allows diversification of Panoptoo systems similar to nucleotide immune signal adaptations in other mechanisms of anti-phage defense^4,6,30,33^. DS-2C protein synthesis of 3′3′-c-di-AMP potentially functions as an additional mechanism to respond to phage infection in CBASS defense, but definition of the specific role of these proteins in bacterial immunity remains an important goal for future analysis.

Joining previous discoveries of nucleotide immune signaling systems in bacterial, plant, and animal antiviral immunity, identification of active DAC domains proteins in Panoptoo defense demonstrates that all known families of cyclic nucleotide signaling enzymes are involved as components in antiviral immunity. Distinct protein domains involved in nucleotide immune signaling now include mcPol / GGDEF / adenylate cyclase ferredoxin-like fold enzymes in type III CRISPR, Panoptes, and Pycsar defense^6,10,11,19,20^; DAC enzymes in Panoptoo defense; CD-NTase and cGLR enzymes in CBASS defense and animal innate immunity^4,5,14,17^; and TIR domains in bacterial, plant, and animal immunity^7,8,13,33,34^. Concomitant with our study, the work of Nabhani and Sullivan et al.^52^ further establishes PanS DAC proteins as synthase enzymes in guard-like anti-phage defense systems. PanS signaling to control PanE activation also expands a growing theme of the role of nucleotide signals as negative regulators in antiviral immunity^18–21^. We find that PanE-like effectors are additionally encoded next to an assortment of proteins of unknown function (Figure 2c), indicating that other yet uncharacterized protein domains may also synthesize nucleotide signals as negative regulators in immunity. Together, our results uncover functions of DAC domains in bacterial antiphage defense and expand our understanding of nucleotide signals that create the chemical language of antiviral immunity.

## Acknowledgements

The authors are grateful to members of the Kranzusch laboratory for helpful comments and discussion. The work was funded by grants to

P.J.K. from the Pew Biomedical Scholars program, the Burroughs Wellcome Fund PATH program, The G. Harold and Leila Y. Mathers Charitable Foundation, the Cancer Research Institute, the Parker Institute for Cancer Immunotherapy, and the National Institutes of Health (1DP2GM146250-01). This paper was typeset with the bioRxiv word template by @Chrelli: www.github.com/chrelli/bioRxiv-word-template

## Author contributions

The study was designed and conceived by A.E.R. and P.J.K. Bioinformatic analysis, biochemical assays, and bacterial growth assays were performed by A.E.R with assistance from D.J.H. Protein purification and structure determination was performed by A.E.R and D.J.H. The figures were prepared by A.E.R. The manuscript was written by A.E.R. and

P.J.K. All authors contributed to editing the manuscript and support the conclusions.

## Materials and Methods

### AlphaFold3 modeling and bioinformatics

AlphaFold3 server^43^ was used to predict the structure of DS-2C and DisA_N as monomers. Five output models were compared in PyMOL to confirm agreement with output model 0 presented in the figures. Models are colored by pLDDT. Pairwise structural comparison between *Ec*DS-2C individual domains, *Ko*DisA_N and protein structures in the PDB was performed using DALI^44^. A Z-score cut-off of 10 for *Ec*DS-2C DAC domain and *Ko*DisA_N was selected to compare relevant homologs while the top 100 hits were plotted for *Ec*DS-2C DACNG, *Ec*DS-2C DACNH, and *Ec*DS-2C C-terminal domain. WebLogos were created by aligning DAC domain and PanS protein sequences using MAFFT (FFT-NS-i iterative refinement method)^53^ and then select regions of the alignment were visualized using WebLogo^3 54^ and colored by chemical property.

S2TMβ protein sequences were downloaded from InterPro (IPR041208), aligned using MAFFT (FFT-NS-i iterative refinement method)^53^, and used to construct a phylogenetic tree in Geneious Prime v2025.2.2 using the neighbor-joining method and Jukes–Cantor genetic distance model with no outgroup. The pie chart in Figure 2 was generated using Prism 11. PanS sequences were obtained through a WebFlags^55^ search of KoDisA_N and 200 sequences encoded next to an S2TMβ effector were aligned using MAFFT (FFT-NS-i iterative refinement method)^53^, and the PanS phylogenetic tree was generated in Geneious Prime v2025.2.2 using the neighbor-joining method and Jukes– Cantor genetic distance model with no outgroup. In each case, manual analysis of sequences were performed to annotate neighboring genes and phylogenetic trees were visualized and annotated in iTOL v7. Genomic neighborhood analysis was performed using WebFlags^55^ and Defense Finder^56^.

### Protein expression and purification

Sequences used in this paper are included in Supplementary Table 1. Diadenylate cyclase proteins were expressed in *E. coli* as previously described^57^. Briefly, expression vectors were transformed into BL21(DE3)RIL (Agilent) plated on MDG media plates (1.5% Bacto agar, 0.5% glucose, 25 mM Na_2_HPO_4_, 25 mM KH_2_PO_4_, 50 mM NH_4_Cl, 5 mM Na_2_SO_4_, 0.25% aspartic acid, 2–50 μM trace metals, 100 μg mL^−1^ ampicillin, 34 μg mL^−1^ chloramphenicol) and grown overnight at 37°C. Five colonies were picked to inoculate 30 mL overnight MDG starter cultures (37°C 230 rpm). 1 L M9ZB expression cultures (47.8 mM Na_2_HPO_4_, 22 mM KH_2_PO_4_, 18.7 mM NH_4_Cl, 85.6 mM NaCl, 1% Cas-Amino acids, 0.5% glycerol, 2 mM MgSO_4_, 2–50 μM trace metals, 100 μg mL^−1^ ampicillin, 34 μg mL^−1^ chloramphenicol) were then inoculated with 10 mL or starting OD_600_ of 0.0475 of MDG starter cultures and then induced with 0.5 mM IPTG after reaching an OD_600_ of ≥1.5 followed by overnight induction (16°C, 230 rpm).

After overnight inductions, cells were centrifuged, re-suspended, and pellets were lysed by sonication in 60 mL lysis buffer (20 mM HEPES pH 7.5, 400 mM NaCl, 10% glycerol, 20 mM imidazole, 1 mM DTT). Lysate was clarified by centrifugation, and resulting supernatants was added to Ni-NTA resin (Qiagen). Resin was washed with lysis buffer, lysis buffer supplemented to 1 M NaCl and then lysis buffer again. Samples were eluted with lysis buffer supplemented to 300 mM imidazole. Samples were then dialyzed overnight in 14 kDa MWCO dialysis tubing (Ward’s Science) with SUMO2-cleavage by hSENP2 as previously described^57^. Proteins were purified by size exclusion chromatography using a 16/600 Superdex 200 column (Cytiva) and stored in gel filtration buffer (20 mM HEPES-KOH pH 7.5, 20 mM KCl, and 1 mM TCEP-KOH). Final proteins were concentrated to >10 mg mL^−1^ using a 30 kDa MWCO centrifugal filter (Millipore Sigma), aliquoted, flash frozen in liquid nitrogen, and stored at −80°C.

### Nucleotide product analysis

Diadenylate cyclase synthesis activity was analyzed by thin-layer chromatography (TLC) as previously described^4,15^. Recombinant protein preparations (5 µM final) were incubated in 10 μL reactions containing 0.5 μL α-^32^P-labeled ATP (approximately 0.4 μCi), 250 μM unlabeled NTPs (ATP, GTP, CTP, UTP) in a final reaction buffer of either 5 mM MgCl_2_, 20 mM Tris-HCl pH 7.5, and 100 mM KCl or 1 mM MnCl_2_, 50 mM Tris-HCl pH 9.0, and 100 mM KCl. Reactions were incubated at 37°C for one hour and subsequently treated with 1 μL Quick CIP phosphatase (New England Biolabs) for 20 min at 37°C to remove unreacted phosphate signal. Each reaction was spotted (0.5 µL) on a 20-cm × 20-cm PEI-cellulose TLC plate. Plates were run with 1.5 M KH_2_PO_4_ solvent until approximately 2.5 cm from the top of the plate, dried at room temperature and exposed to a phosphor-screen before signal detection with a Typhoon Trio Variable Mode Imager System (GE Healthcare). TLC images were adjusted for contrast using FIJI^58^ and quantified using ImageQuant (8.2.0).

Nuclease P1 cleavage analysis was performed using PanS reactions labeled with either α-^32^P-labeled ATP or α-^32^P-labeled UTP as previously described^4,15,50^. Briefly, radiolabeled nucleotide products were incubated with nuclease P1 (80 mU; N8630, Sigma) in buffer (30 mM NaOAc pH 5.3, 5 mM ZnSO_4_, and 50 mM NaCl) for 30 min followed by 30 min of treatment with Quick CIP (NEB).

### Liquid chromatography-mass spectrometry

Enzymatic synthesis of DAC products for LC-MS analysis was performed using reactions containing 5 µM DAC enzyme, 200 µM ATP, 200 µM GTP, 200 µM CTP, 200 µM UTP, 100 mM KCl, 1 mM DTT, 5 mM MgCl_2_, 1 mM MnCl_2_, and 20 mM Tris-HCl at pH 7.5 or 9.0. Reactions were incubated at 37°C for 1 hour and the nucleotide product was recovered after boiling reactions at 95°C for 5 min followed by filtering reactions through a 3 kDa MWCO filter (Amicon) to remove protein. Co-purified PanS nucleotide products were recovered from purified PanS enzymes after boiling at 95°C for 5 min followed by filtering through a 3 kDa MWCO filter (Amicon). Nucleotide products were separated on an Agilent 1260 HPLC equipped with a diode array detector and an Agilent 6125 single quadrupole mass spectrometer in negative ion mode using a reverse-phase Agilent InfinitiLab Poroshell SB-Aq column (2.7-µm particle size, 2.1-mm inner diameter, 100-mm length) at a flow rate of 0.45 ml min^−1^ with a gradient from 100% solvent A (0.1% ammonium formate) to 100% solvent B (methanol). Data were collected by OpenLAB CDS v.2.7 software and are representative of at least two independent experiments.

### Bacterial growth assays

Panoptoo operons or PanE proteins alone were synthesized and cloned into an arabinose-inducible pBAD vector^23^ (Twist). *E. coli* BW25113 Δ9 (NCTC 14365)^59^ cells were transformed with Panoptoo plasmids and plated onto LB supplemented with 1% glucose and 100 µg mL^−1^ ampicillin. One colony from each transformation was picked into 3 mL LB with 1% glucose and 100 µg mL^−1^ ampicillin and grown overnight at 37°C with shaking at 230 rpm. Cells were pelleted, re-suspended in PBS (137 mM NaCl, 2.7 mM KCl, 10 mM Na_2_HPO_4_, 1.8 mM KH_2_PO_4_), and serially diluted in a 96-well plate. Five µL of each serial dilution was spotted onto LB plates supplemented with 100 µg mL^−1^ ampicillin and 0.02% L-arabinose as well as LB plates supplemented with 100 µg mL^−1^ ampicillin and 1% glucose. Plates were allowed to dry for 1 hour and incubated overnight at 37°C.

For phage sponge co-expression experiments, pBAD-Panoptoo plasmids were co-transformed with sfGFP, SPO1 Tad2, or SPO1 Acb4 proteins encoded on an IPTG-inducible pCA24N vector into *E. coli* BL21 (DE3) (NEB) cells and plated onto LB supplemented with 1% glucose and 100 µg mL^−1^ ampicillin. One colony from each transformation was picked into 3 mL LB with 1% glucose and 100 µg mL^−1^ ampicillin and grown overnight at 37°C with shaking at 230 rpm. Cells were pelleted, re-suspended in PBS (137 mM NaCl, 2.7 mM KCl, 10 mM Na_2_HPO_4_, 1.8 mM KH_2_PO_4_), and serially diluted in a 96-well plate. Five µL of each serial dilution was spotted onto LB plates supplemented with 100 µg mL^−1^ ampicillin and 0.02% L-arabinose and 0.1 µM IPTG as well as LB plates supplemented with 100 µg mL^−1^ ampicillin and 1% glucose. Plates were allowed to dry for 1 hour and incubated overnight at 37°C.

### Protein crystallization and structural determination

Crystals of native *Re*PanS were grown for three days in reservoir solution (0.1 M HEPES pH 7.5, 20% PEG 8000) at 18°C using hanging-drop vapor diffusion before cryoprotection with the reservoir solution supplemented with 20% ethylene glycol and freezing in liquid nitrogen. X-ray diffraction data were collected at Advanced Photon Source beamline 24-ID-E. Data were processed with XDS and Aimless^60^. Experimental phase information was determined by molecular replacement using an AlphaFold3 predicted structure in PHENIX^61^. Model building was completed in Coot^62^ and then refined in PHENIX. Statistics were analyzed as presented in Extended Data Table 1 and structure figures were produced in PyMOL. The final structure was refined to stereochemistry statistics for Ramachandran plot (favored/allowed), rotamer outliers and MolProbity score as follows: 98.02%/1.80%, 0.95% and 1.28.

---

**Extended Data Figure 1.**
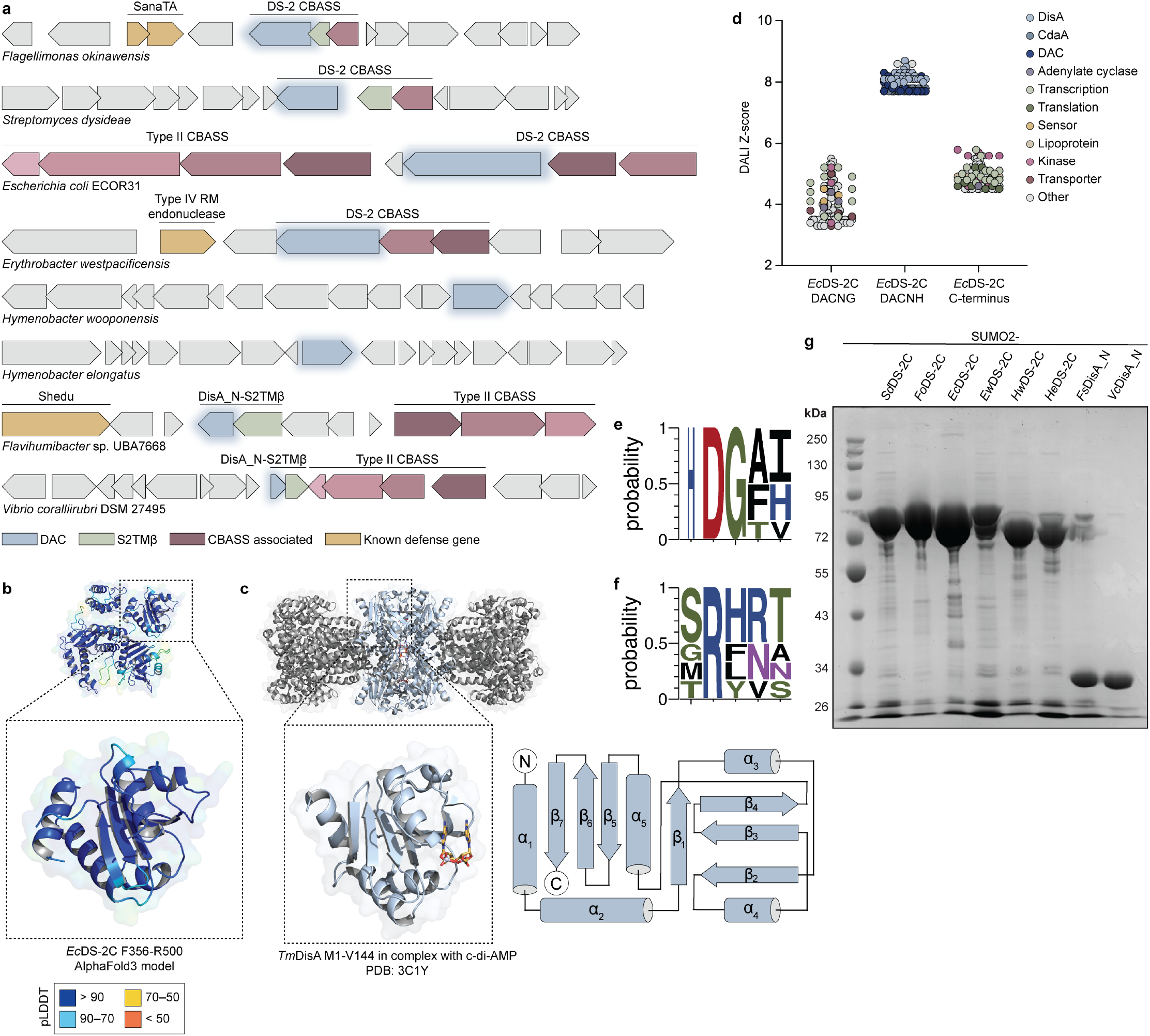
DAC domains located in defense islands share homology with characterized diadenylate cyclases. **a**, Genomic neighborhood analysis for selected DS-2C and DisA_N proteins. **b**, AlphaFold3 predicted structure model of the full-length *Ec*DS-2C protein colored by pLDDT. **c**, Structure of the canonical DAC domain protein DNA integrity scanning protein A (DisA) from T. *maritima* (PDB: 3C1Y) with the DAC domain highlighted in blue. Below: zoom-in of the *Tm*DisA N-terminal DAC domain. Right: topology diagram for the *Tm*DisA DAC domain. **d**, Z-score structural similarity plot demonstrating homology between model predictions of the individual domains of *Ec*DS-2C with representative structures in the Protein Data Bank. The top 100 hits are shown. **e–f**, WebLogo showing the conservation of DAC catalytic residues for two representative DisA sequences, two representative DS-2C sequences, and two representative DisA_N sequences. **g**, Coomassie-stained 10% SDS-PAGE analysis of purified DS-2C and DisA_N proteins.

**Extended Data Figure 2.**
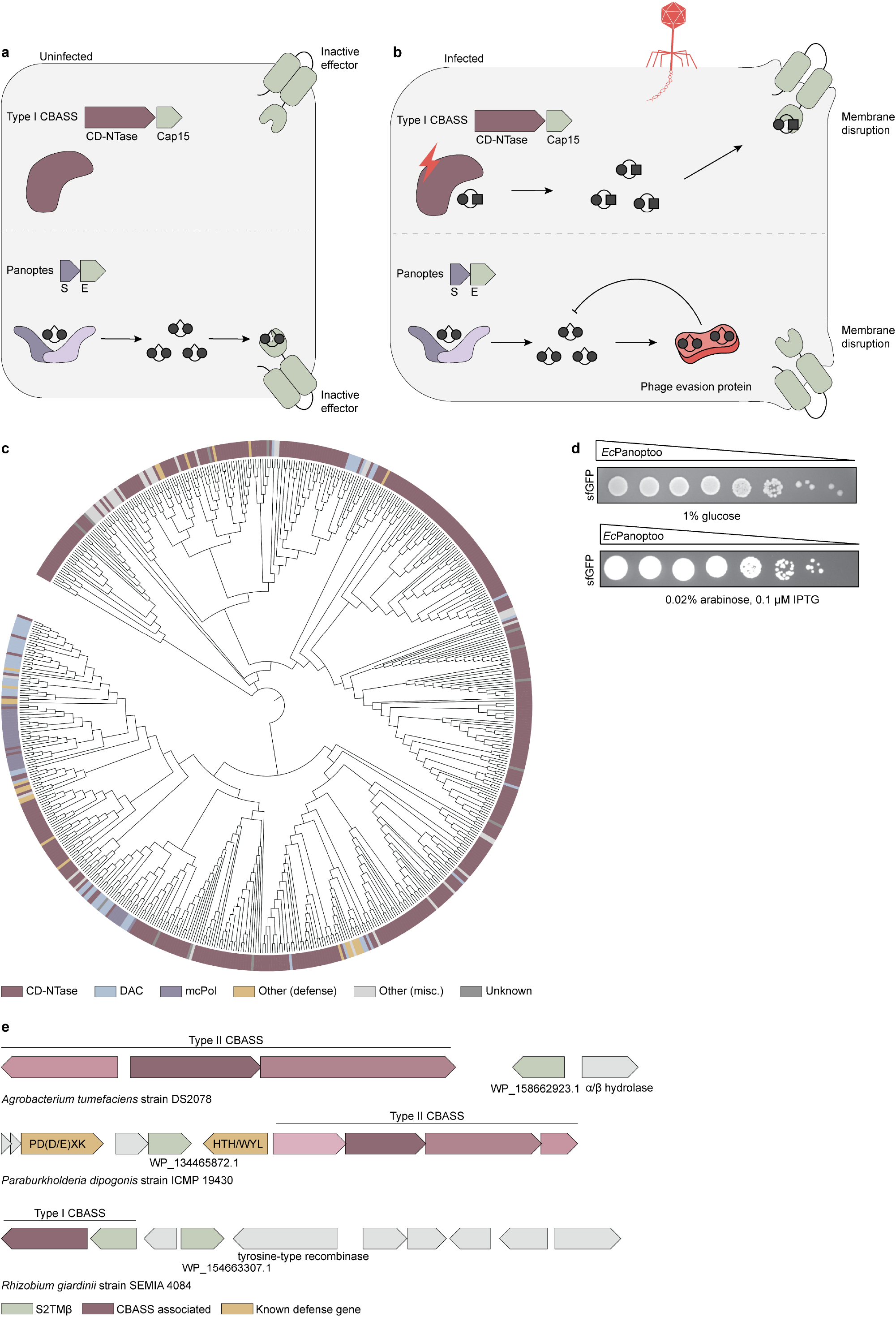
S2TMβ effectors are encoded next to diverse nucleotide signal generating enzymes. **a–b**, Model schematic of the function of additional examples of S2TMβ effectors in CBASS and Panoptes anti-phage defense. In type I CBASS, CD-NTase nucleotide immune signal synthesis is inactive in the absence of phage infection. Upon phage infection, the CBASS CD-NTase enzyme synthesizes a nucleotide signal that then activates a downstream S2TMβ effector protein to induce membrane disruption and an abortive infection response that limits phage propagation. In Panoptes, OptS nucleotide signal synthesis is constitutive and generates an immune signal that negatively regulates OptE effector function. Upon phage infection, if the phage encodes an anti-CBASS immune evasion protein that disrupts nucleotide signaling, repression of OptE is released to induce membrane disruption and an abortive infection response that limits phage propagation. **c**, Phylogenetic analysis of 617 S2TMβ sequences (InterPro IPR041208) annotated by the neighboring gene. **d**, Bacterial growth assay of the co-expression of *Ec*Panoptoo from an arabinose inducible pBAD plasmid with sfGFP from an IPTG inducible vector in *E. coli*. Data shown are representative of three independent experiments. **e**, Genomic neighborhood analysis of S2TMβ effectors encoded next to uncharacterized proteins highlights additional potential classes of nucleotide signal generating enzymes.

**Extended Data Figure 3.**
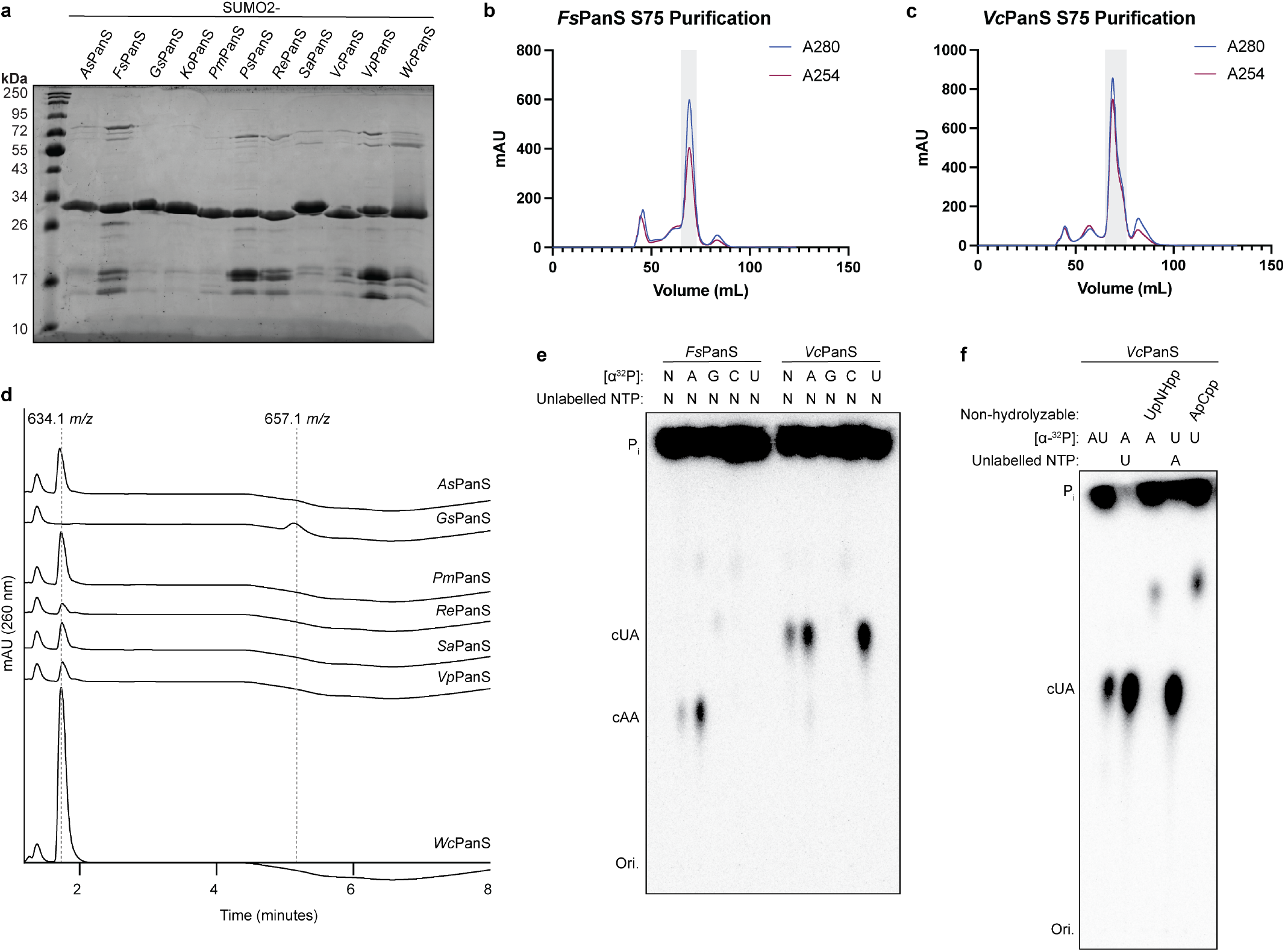
Biochemical characterization of c-di-AMP and cUA synthesizing PanS enzymes. **a**, Coomassie-stained 15% SDS-PAGE of purified PanS enzymes. **b–c**, Size-exclusion chromatogram (16/600 S75) for *Fs*PanS (b) and *Vc*PanS (c). Shading indicates which peak was used for downstream analysis. **d**, LC-MS analysis of reactions performed at pH 9.0 with MgCl_2_ and MnCl_2_ verifies that *Gs*PanS synthesize c-di-AMP while all other homologs synthesize cUA. **e**, Biochemical deconvolution of PanS reactions after incubation with α^32^P-labeled and unlabeled NTPs. For *Fs*PanS, a product is only visualized after incubation with labeled ATP, confirming the cyclized product contains only adenosine. For *Vc*PanS, a product is visualized after incubation with labeled ATP or UTP, confirming the cyclized product contains adenosine and uridine. **f**, Analysis of *Vc*PanS reactions with pairwise combination of α^32^P-labeled and non-hydrolyzable NTPs reveals there is no preference for the order of bond formation during 3′3′-cUA synthesis. Data shown in **d–f** are representative of three independent experiments.

**Extended Data Figure 4.**
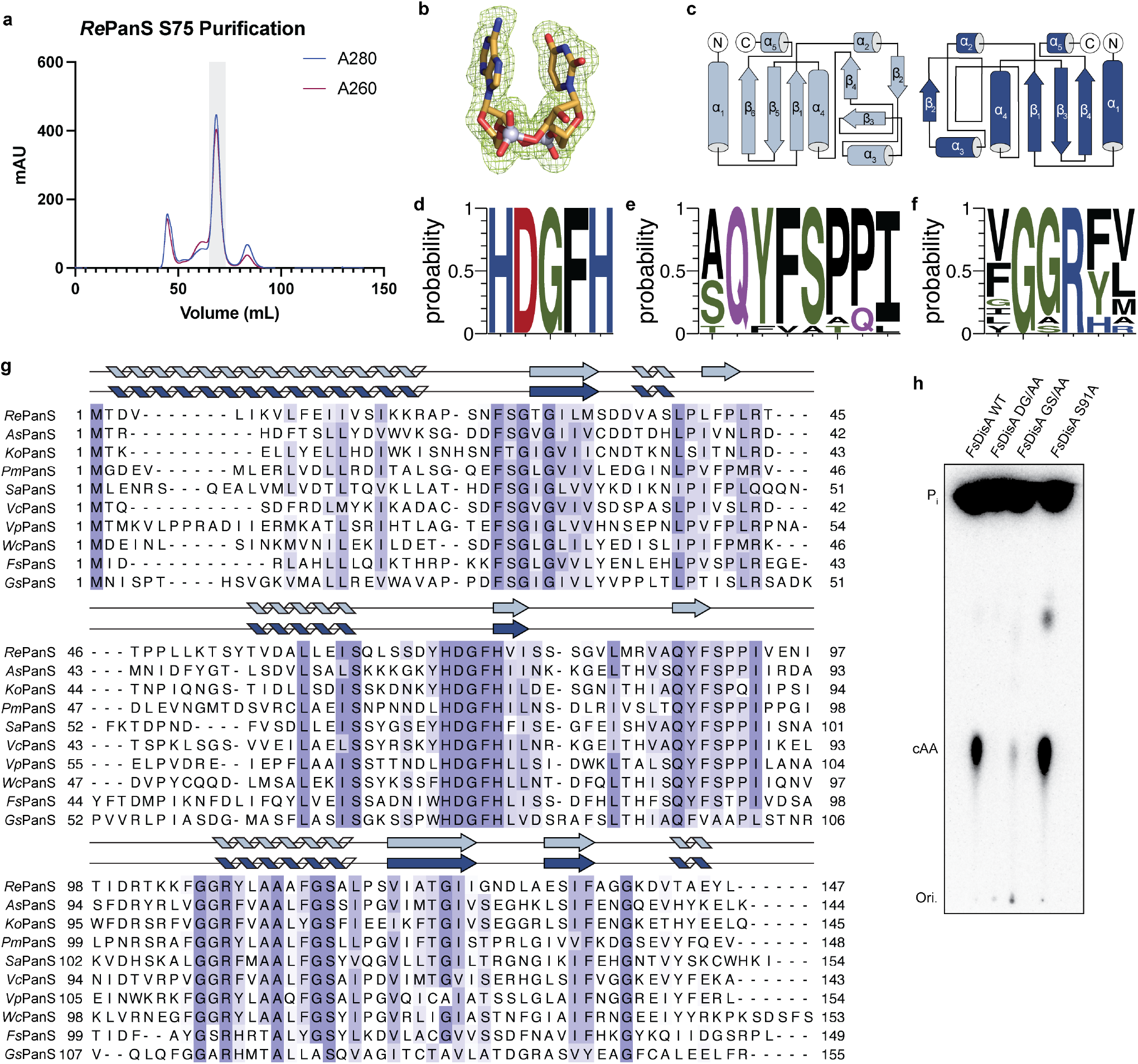
Structural analysis of PanS. **a**, Size-exclusion chromatogram (16/600 S75) for *Re*PanS. Shading indicates the protein peak selected for downstream analysis. **b**, F_O_–F_C_ polder omit map (σ = 3) (green mesh) of the 3′3′-cUA ligand in the *Re*PanS–3′3′-cUA structure. **c**, Topology diagram for the two protomers of *Re*PanS with the adenine-contacting protomer colored in light blue and the uracil-contacting protomer colored in dark blue demonstrates the change in the loop conformation that allows for synthesis of the asymmetric signaling product. **d–f**, Analysis of PanS homologs demonstrates conservation of catalytic residues DGF (**d**) and GGR (**f**) as well as residues in the active site lid loop responsible for nucleobase selection (**e**). **g**, Structure guided sequence alignment of PanS homologs colored by BLOSUM62 score. **h**, Mutations in the *Fs*PanS conserved catalytic active site residues (DGF and GGR) result in loss of cAA production, while residue S91 is dispensable for product formation. Data are representative of three independent experiments.

